# Haplotype-resolved genetic mapping using selfed progeny enables simultaneous genome assembly validation and trait dissection in a heterozygous plant

**DOI:** 10.64898/2026.10.22.760661

**Authors:** Insu Lim, Sae Hyun Lee, Woohyeon Cho, Junhyun Kim, Jee Young Park, Murukarthick Jayakodi, Tae-jin Yang

## Abstract

Genetic linkage maps are essential for genome analysis and molecular breeding, but developing suitable mapping populations remains challenging in many plant species. Approaches requiring inbred parental lines are particularly difficult to implement in plants with long generation times or severe inbreeding depression. We developed haplotype-resolved genetic mapping (HRGM) to construct linkage maps from a single heterozygous individual and its selfed progeny without requiring inbred parental lines. HRGM combines haplotype-resolved unitig assemblies from the S_0_ parent with genotype data from its S_1_ progeny, using sequence differences between the two haplotypes to define phased SNP markers. Application to Cynanchum wilfordii generated a high-density genetic map comprising 8,147 SNP markers consolidated into 464 bins across 11 linkage groups, spanning 786.4 cM. The map enabled the detection of two loci associated with root development. Recombination patterns derived from the S_1_ population showed strong concordance with the chromosome-scale genome assembly and exhibited recombination suppression in putative centromeric regions. HRGM integrates recombination-based genome assembly validation and trait mapping within a single framework. By using one generation of selfed progeny and obtaining linkage-phase information directly from phased assemblies, it simplifies mapping population development and extends the utility of genomic resources for genetic analysis and molecular breeding, particularly in species where selfed progeny can be obtained but conventional mapping populations are difficult to establish.

## Introduction

Genetic linkage maps have long served as the primary foundation for interpreting genome structure, enabling the ordering of markers through meiotic recombination and supporting trait mapping (Jaganathan *et al*., 2020), positional cloning (Schneeberger, 2014), and genome assembly (Fierst, 2015). In early plant genomics, including *Arabidopsis thaliana* and rice (*Oryza sativa*) (Sasaki, 2005; The Arabidopsis Genome Initiative, 2000), linkage maps were indispensable for anchoring sequence contigs and resolving chromosome-scale organization. However, the role of linkage maps has not diminished with the advent of modern sequencing technologies; rather, they remain the only direct experimental measure of recombination and thus a critical benchmark for validating genome assemblies (Cui *et al*., 2022; Kong *et al*., 2023). Recent advances in long-read sequencing and chromatin conformation capture (Hi-C) have transformed genome assembly, making chromosome-scale, haplotype-resolved assemblies increasingly accessible across diverse plant lineages. Thousands of plant genomes are now available, including those from medicinal, woody (Cheng *et al*., 2025; Michael, 2026), and previously underrepresented plant species (Song *et al*., 2024; Yang *et al*., 2021). Pangenome studies further extend these resources by capturing structural variation and haplotype diversity at population scale (Guo *et al*., 2025; Shi *et al*., 2024). Yet, despite this rapid expansion, most assemblies remain validated primarily by computational consistency rather than independent genetic evidence. As a result, assigning highly similar homologous sequences to their correct chromosomal locations and distinguishing true structural variation such as inversions, translocations and presence/absence variations, from assembly artifacts remains challenging, particularly in highly repetitive and heterozygous genomes (Kong *et al*., 2023; Mahmoud *et al*., 2019; Tørresen *et al*., 2019; Whibley *et al*., 2021). These limitations reduce accuracy in downstream analyses such as molecular marker development and quantitative trait loci (QTL) mapping.

This disconnect reflects a deeper limitation in current genetic mapping frameworks. Conventional linkage mapping relies on segregating populations derived from crosses between inbred parental lines, an approach that is incompatible with many important crop plant species characterized by long generation times (Migicovsky and Myles, 2017; van Nocker and Gardiner, 2014; Roeder and Jill Harrison, 2019), high heterozygosity, and inbreeding depression (Charlesworth and Willis, 2009; Zhang *et al*., 2019). Alternative strategies, including pseudo-testcross designs in F₁ populations, partially address this constraint but typically yield fragmented, parent-specific maps and fail to fully exploit genome-wide marker information (Maliepaard *et al*., 1997; Ooijen, 2011). Selfed (S_1_)-based approaches offer a potential route to mapping without crossing, but their utility has been limited by incomplete linkage-phase information and pervasive segregation distortion (Di Pierro *et al*., 2016; Tian *et al*., 2015; Ward *et al*., 2005). At the same time, haplotype-resolved genome assemblies have fundamentally changed the information landscape. They explicitly encode the two parental chromosome sets within a single individual, capturing phased sequence variation at genome scale (Guk *et al*., 2022; Zhang *et al*., 2021). Despite this, current mapping approaches do not leverage haplotype-resolved assemblies to infer linkage directly, leaving a gap between genome assembly and genetic mapping. Here, we introduce haplotype-resolved genetic mapping (HRGM), a framework that integrates haplotype-resolved unitig assemblies with S_1_ population data to construct linkage maps without the need for a biparental population derived from homozygous parental lines. HRGM infers linkage phase directly from a single heterozygous individual, converting phased sequence variation into a dense set of informative markers that can be unambiguously genotyped across progeny. By aligning recombination patterns with haplotype structure, this approach enables simultaneous genetic mapping and genome assembly validation within a unified pipeline.

We apply HRGM to *Cynanchum wilfordii* (henceforth CW) (2n=2x=22), a perennial medicinal root crop with high heterozygosity, generating a high-density linkage map and identifying QTL associated with root development. More broadly, HRGM establishes a generalizable strategy for exploiting haplotype-resolved assemblies in species where conventional mapping populations are impractical, bridging a critical gap between genome assembly and functional genetics.

## Results

### Conceptual Framework and Overview of the HRGM-based Genetic Mapping

In conventional linkage mapping, genetically differentiated parental lines are typically crossed to generate segregating populations. However, the development of inbred parental lines or advanced mapping populations may require multiple generations and can be particularly difficult in highly heterozygous species and crops with long generation times (Fig. 1A). In contrast, haplotype-resolved genetic mapping (HRGM) circumvents this limitation by exploiting the phased contiguous sequences of a single heterozygous individual (S_0_) and its selfed progeny (S_1_) to construct an integrated linkage map without inbred parents and with minimal inbreeding depression (Fig. 1B). This pipeline transforms within-individual heterozygosity into a dense and fully phased marker system, which facilitates the rapid construction of high-resolution genetic maps without requiring chromosome-scale haplotype-resolved assemblies or inbred parental lines (Fig. 2A).

**Figure 1.**
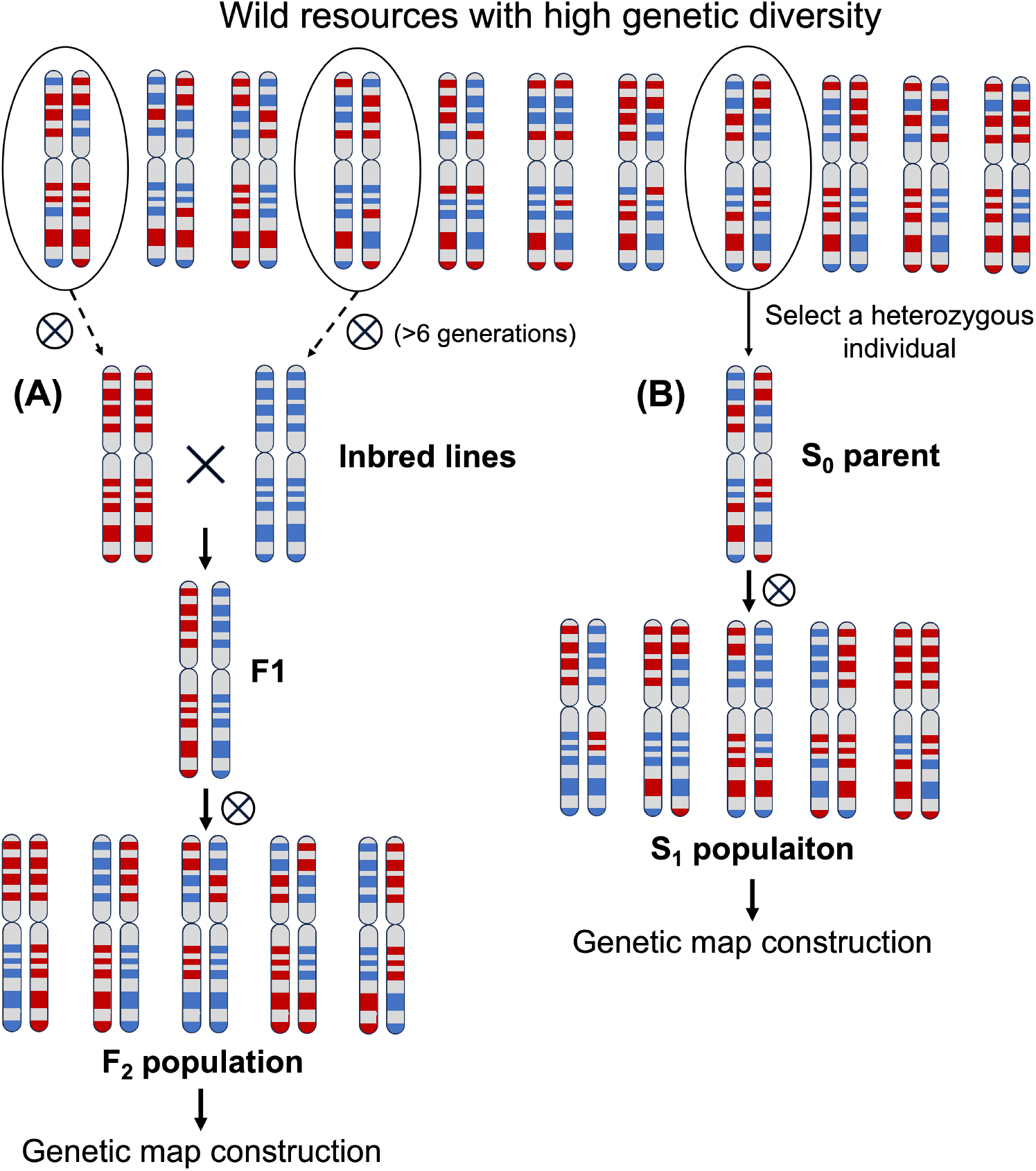
Comparison of two mapping strategies using a wild plant population. **(A) Traditional approach to construct a mapping population.** Two parental inbred lines are developed through repeated selfing for six to eight or more generations, crossed to produce an F₁ hybrid, and subsequently selfed to generate an F₂ mapping population. The process requires more than eight generations. **(B) The haplotype-resolved genetic mapping developed in this study**. A highly heterozygous plant can be used directly for haplotype-resolved genome assembly and self-pollinated to generate S_1_ progeny for use as a mapping population.

**Figure 2.**
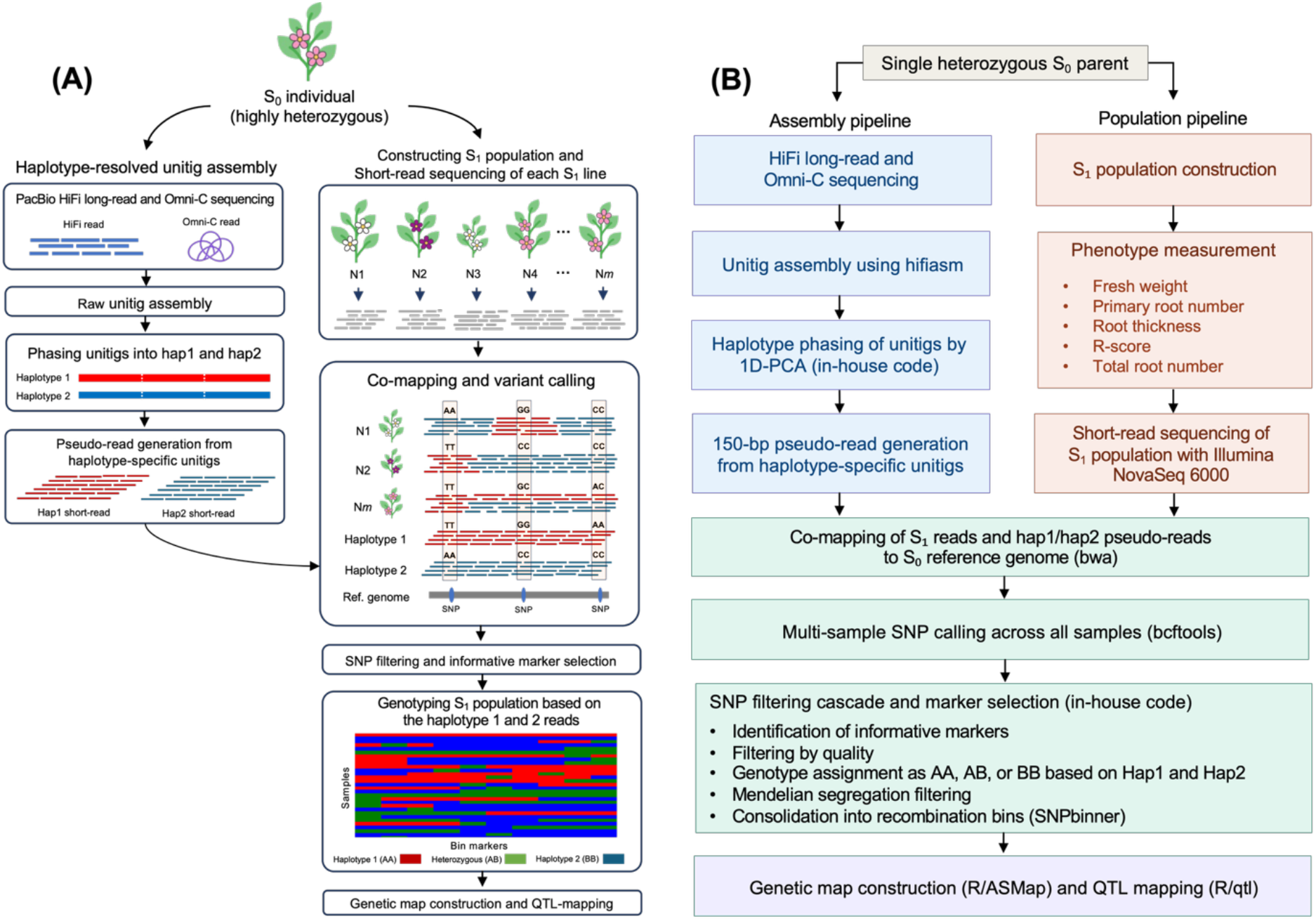
Conceptual framework and pipeline of haplotype-resolved genetic mapping (HRGM). (A) Schematic overview of the HRGM workflow. HiFi reads from the S_0_ individual are assembled into unitigs, phased into hap1 and hap2 using Omni-C contact information, and fragmented into artificial 150-bp pseudo-reads. An S_1_ segregating population is developed from the S_0_ parent, and Illumina short reads are produced for each S_1_ individual. The pseudo-reads and S_1_ reads are co-mapped to a reference genome for SNP calling. Informative SNPs fixed within each haplotype and polymorphic between them are genotyped as AA, AB, or BB, filtered for Mendelian segregation, and merged into bin markers for genetic map construction. (**B) Corresponding bioinformatic implementation of the HRGM pipeline.** The pipeline proceeds along two parallel tracks, assembly (blue) and population (pink), that converge at joint genotyping (green), encompassing co-mapping, SNP calling, and marker selection. The resulting bin markers are used for genetic map construction and QTL mapping (purple). Key bioinformatic tools and in-house code used at each step are indicated.

The HRGM pipeline proceeds through a series of assembly, phasing, genotyping, marker selection, and genetic map construction steps (Fig. 2B). In the assembly step, HiFi long-read and Omni-C sequencing data are generated from the S_0_ individual. The HiFi reads are assembled into unitigs using hifiasm (Cheng *et al*., 2021), and these unitigs are then phased into two haplotype sets, hap1 and hap2, by applying one-dimensional principal component analysis (1D-PCA) to the Omni-C contact matrix using an in-house pipeline. Each haplotype set represents one of the two homologous chromosome sets of the S_0_ individual and provides haplotype-specific sequence information for genotyping the S_1_ progeny. The phased unitigs are subsequently decomposed into artificial 150-bp reads using a sliding-window approach, generating haplotype-specific pseudo-reads for joint SNP calling.

In parallel, the S_0_ individual is selfed to generate an S_1_ mapping population, and each S_1_ individual is phenotyped for traits of interest and subjected to whole-genome short-read sequencing (Fig. 2B). S_1_ reads and haplotype-specific pseudo-reads are then co-mapped to the S_0_ reference genome for multi-sample SNP calling. The SNPs that are fixed within each haplotype but polymorphic between haplotypes are retained as informative markers. Based on these haplotype-diagnostic SNPs, each S_1_ individual is assigned an unambiguous genotype at each locus as AA for hap1 homozygotes, AB for heterozygotes, and BB for hap2 homozygotes (Fig. 2A, B). The markers are consolidated into recombination bins and assembled into linkage groups, yielding a high-density linkage map suitable for QTL analysis and genome assembly validation (Fig. 2B).

### Phased Unitig Assembly and Genotyping of S_1_ Population

To demonstrate our concept, we generated haplotype-resolved unitig assemblies for *C. wilfordii* (accession 9p) using previously produced HiFi long reads and Omni-C. The assembly comprised 8,115 raw unitigs (N50 = 113.65 kb; total length = 485.16 Mb). A total of 1,339 unitigs with Omni-C reads were obtained, and the 7 homozygous unitigs with mapping coverage >10× were removed (Fig. 3A). The remaining unitigs were then grouped into two balanced haplotypes (hap1: 657 unitigs, 125.7 Mb; hap2: 675 unitigs, 126.5 Mb) (Fig. 3B and Table S_1_), providing sequence representations of the two homologous haplotypes of the S_0_ individual.

**Figure 3.**
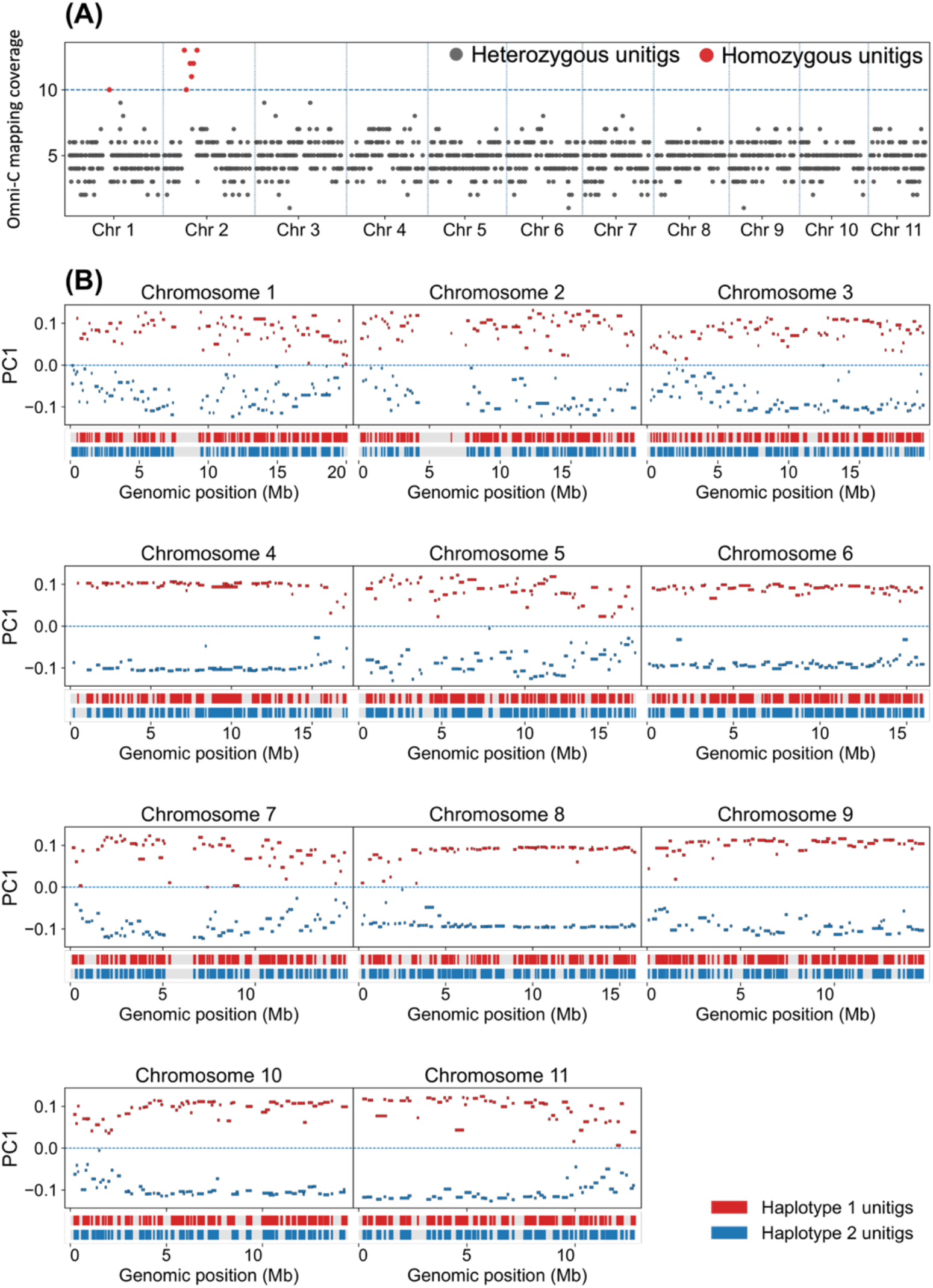
Unitig Filtering and Phasing. **(A)** Omni-C mapping coverage on unitigs across the 11 chromosomes. Red and gray dots indicate homozygous and heterozygous unitigs with coverage >10× and ≤10×, respectively. **(B)** The result of one-dimensional PCA for haplotype phasing. The PC1 value was calculated using Omni-C contact matrices. The red and blue lines are unitigs for haplotypes 1 and 2, respectively. Below each panel, phased regions are illustrated as two horizontal tracks representing haplotype 1 and haplotype 2.

To convert phased assemblies into genetic markers, haplotype-specific unitigs were decomposed into artificial 150-bp reads using a sliding-window approach with a step size of 3 bp. This process generated ∼42 million pseudo-reads per haplotype (Table 1). These pseudo-reads served as phase-defined references (hap1 and hap2), which enabled direct genotyping of S_1_ individuals. Whole-genome resequencing of 52 S_1_ progeny yielded an average of 19.1 million reads per individual (∼15-fold coverage) (Table 1). Joint mapping of S_1_ genomic reads together with hap1-and hap2-derived pseudo-reads to the *C. wilfordii* reference genome identified 3.96 million SNPs. The haplotype-derived pseudo-reads served as phase-defined references that allowed each SNP allele to be assigned to its haplotype of origin. Using these pseudo-read genotypes, the HRGM pipeline retained only SNPs that were homozygous and fixed within each haplotype pseudo-read set but carried alternative alleles between hap1 and hap2. To reduce false-positive diagnostic SNPs caused by ambiguous mapping of artificial short reads, candidate markers were further validated through direct alignment of the corresponding haplotype unitigs to the reference genome. Only SNPs with exact and unambiguous nucleotide support from both haplotype 1 and haplotype 2 unitig alignments were retained. This procedure identified 53,095 haplotype-diagnostic SNPs (Table 1).

**Table 1.**
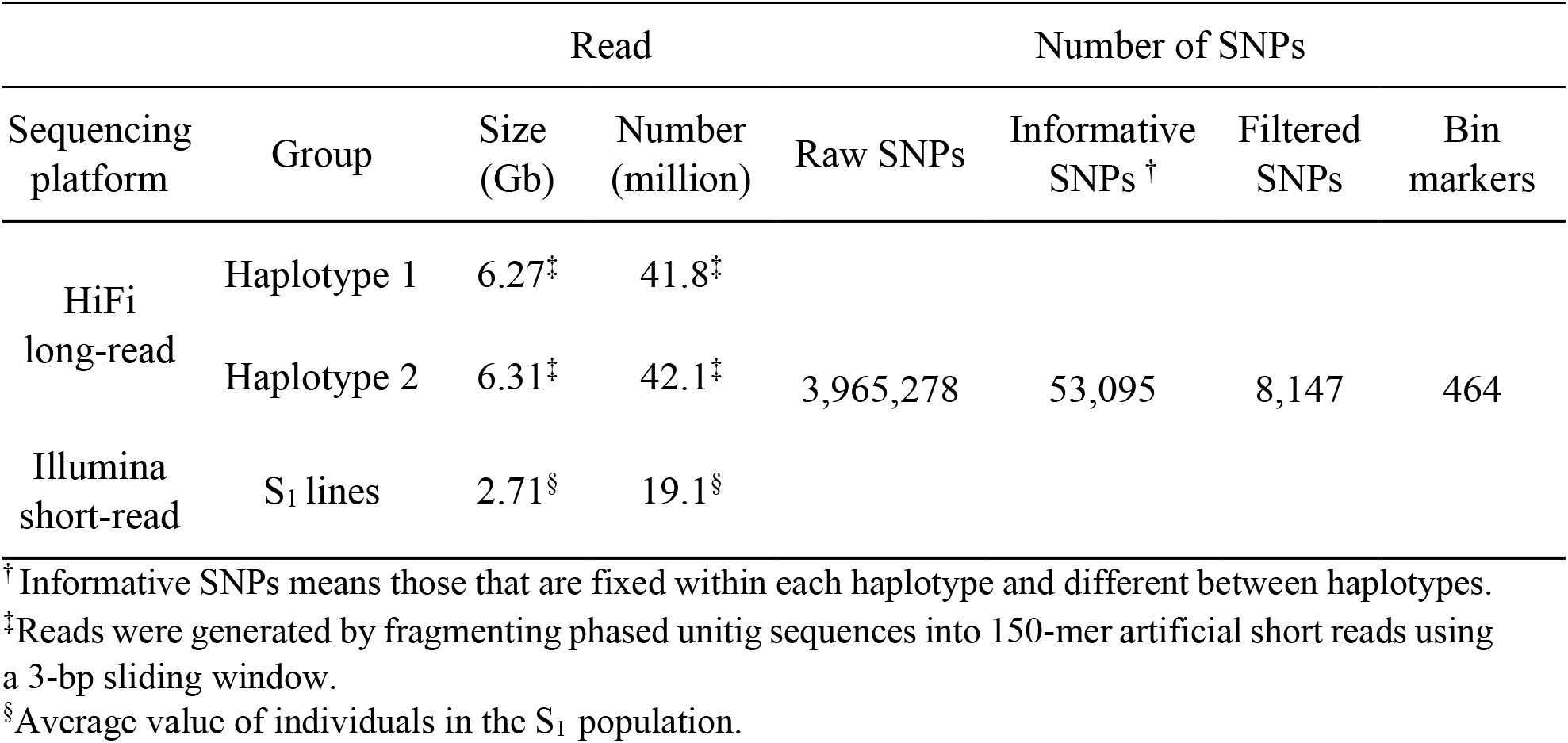
Sequencing and SNP calling information.

These SNPs were distributed across all chromosomes, providing genome-wide coverage suitable for linkage analysis (Figure S1). Following stringent filtering for genotype quality, sequencing depth, and Mendelian segregation (1:2:1), 8,147 high-confidence SNP markers were retained. Genotypes were unambiguously assigned as AA, AB, or BB according to their correspondence with hap1-and hap2-specific alleles, and redundant markers were consolidated into 464 recombination bins. To assess potential mapping artifacts arising from structural differences between haplotypes, we examined synteny between hap1 and hap2. The analysis revealed only localized, small-scale rearrangements with no broad-scale inversions or translocations (Figure S2). Consistently, segregation distortion across the 464 bin markers was minimal and not concentrated in any particular chromosomal region.

### Construction of Linkage-Based Genetic Map

Using 464 bin markers, we constructed 11 linkage groups corresponding to the 11 chromosomes (Fig. 4A and Table S2). The genetic map spanned 786.4 cM, with linkage group lengths ranging from 28.2 to 127.2 cM and a mean inter-marker interval of 1.74 cM (Table S2). The marker order was highly concordant between the physical and genetic maps across all 11 linkage groups, with no large-scale discordance in marker order observed (Fig. 4A). Recombination patterns showed elevated recombination in distal chromosomal regions and suppression in putative centromeric regions (Fig. 4B), consistent with canonical patterns observed in other plant genomes (Brazier and Glémin, 2022). Together, these results indicate that HRGM captures biologically meaningful recombination patterns while maintaining structural agreement with the physical genome.

**Figure 4.**
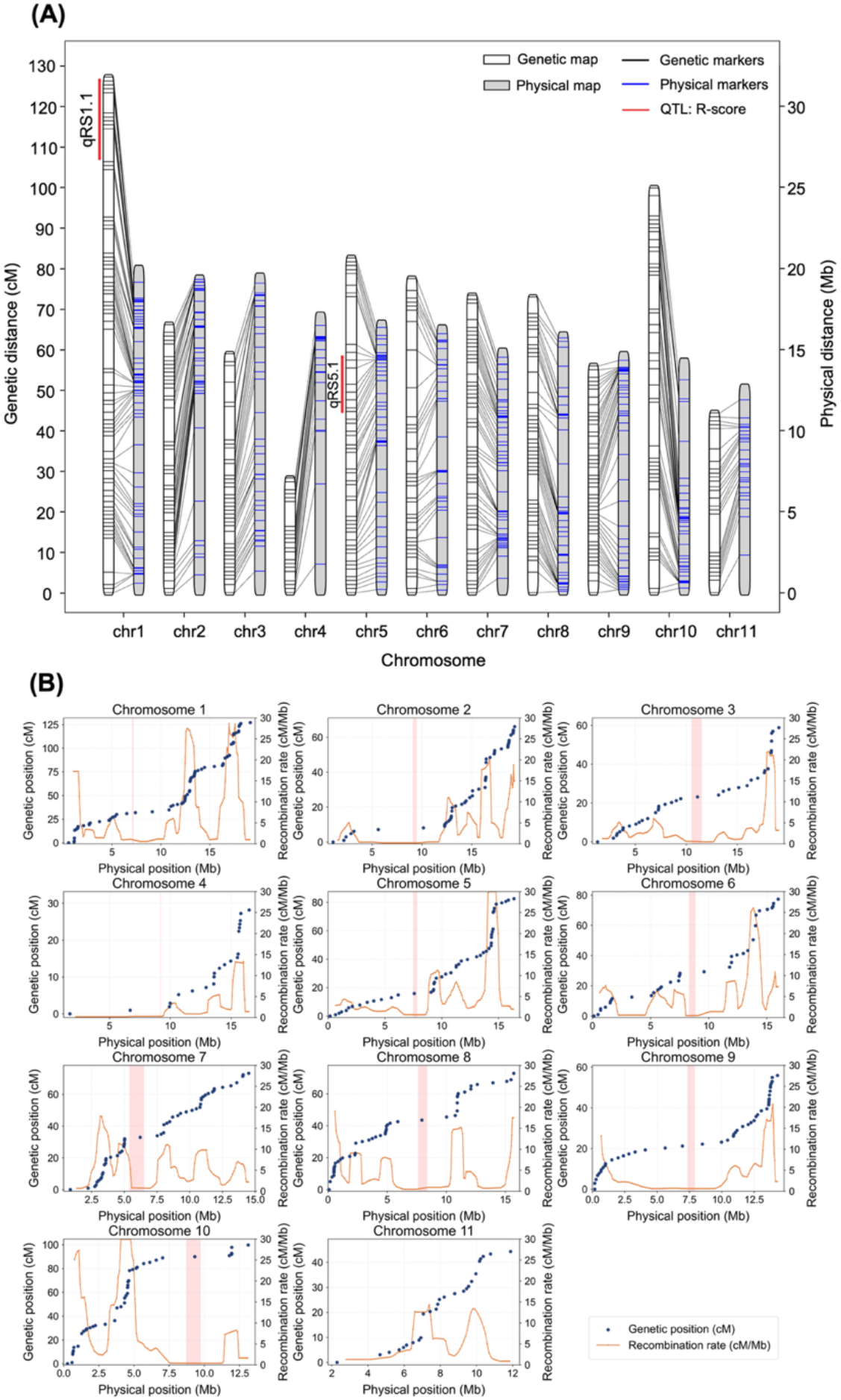
Genetic map construction. (A) Genetic map and corresponding physical map. The linkage map distance (cM) and the corresponding pseudomolecule length (Mb) are shown on the left and right y-axis, respectively. The black and blue lines indicate bin markers on the genetic and physical map, respectively. The bin markers are connected to show a linear order between linkage groups and pseudomolecules. The identified QTLs were anchored to their linkage map positions and were marked in red color. **(B) Correlation between genetic distance and physical distance.** The left and right y-axes indicate genetic distance (cM) and recombination rate (cM/Mb), respectively, and the x-axis indicates physical position (Mb). Blue dots represent individual bin markers, and red boxes indicate the putative centromeric region of each chromosome. Orange dotted lines represent recombination rates calculated using a 0.1-Mb sliding window along each chromosome.

To assess whether the four linkage-map intervals exceeding 10 cM were caused by haplotype-switching errors, we performed genotype-state swap tests for the bins associated with these intervals. The AA and BB genotype states of each target bin were reciprocally exchanged across the S_1_ population, and the linkage map was reconstructed. The four intervals were located on chromosomes 1, 5, and 10, including two intervals on chromosome 10 (Fig. 5 and Figure S3). In all four tests, the exchanged genotype patterns showed abrupt discontinuities relative to the flanking markers, and the modified bins separated from their original linkage groups.

**Figure 5.**
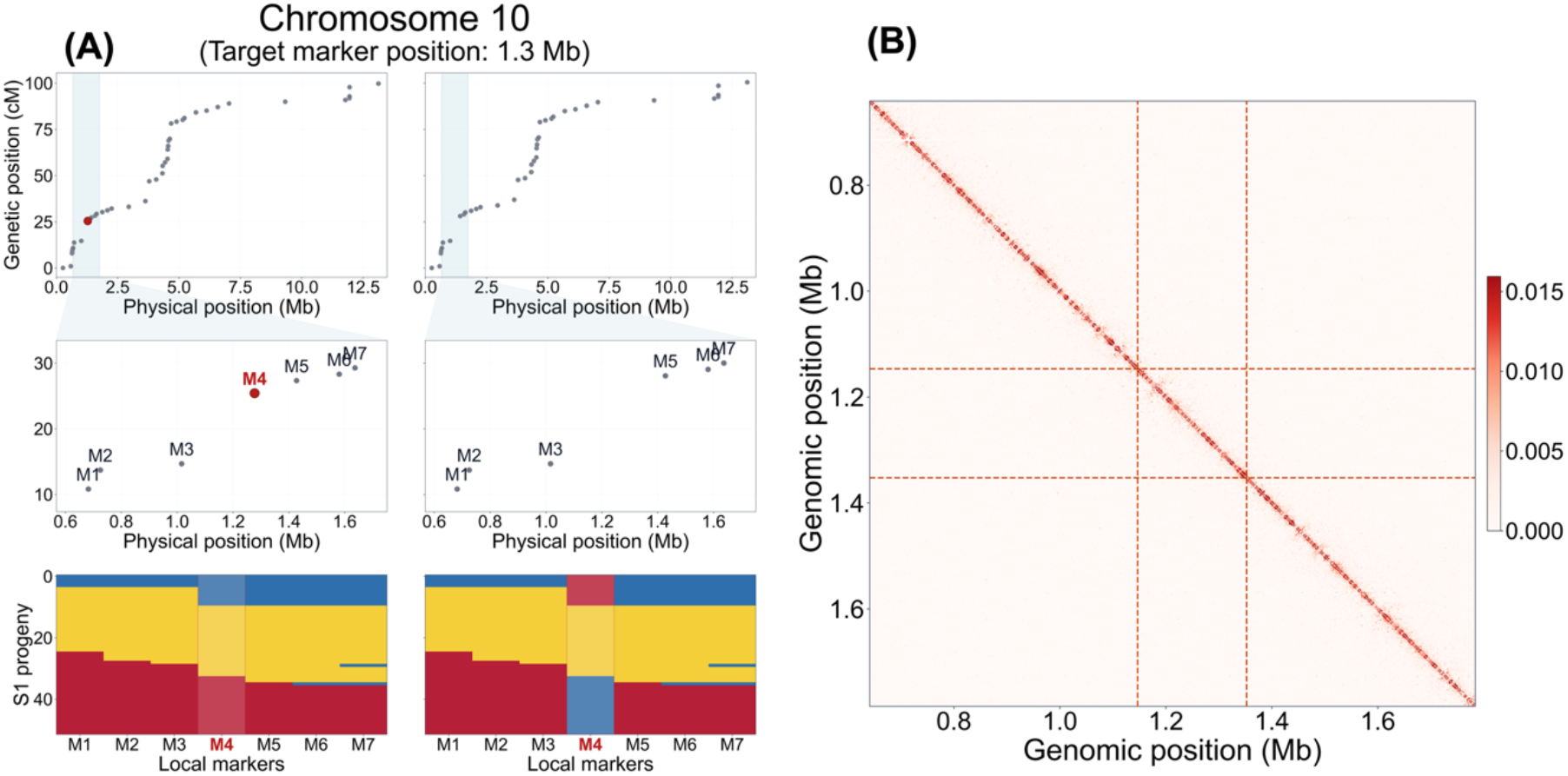
Evaluation of haplotype-phase consistency using an A/B genotype-state swap test. (A) Comparison of the original and reconstructed linkage maps for the target bin chr10-1277289, at which an abrupt increase in genetic distance exceeding 10 cM was observed. The left panels show the original genetic map and S_1_ bin-genotype pattern, whereas the right panels show those reconstructed after reciprocal exchange of the A and B genotype states of the target bin. The target bin is shown in red. (B) Local Omni-C contact map of the target region. Orange dashed lines indicate the boundaries of the target bin, and the color scale represents the Omni-C contact signal.

In contrast, although the original genotype patterns contained several recombination transitions near these regions, these transitions remained consistent with the broader patterns of adjacent markers and supported the original bin assignments (Fig. 5 and Figure S3). Local Omni-C contact maps based on the reference genome also showed no evidence of structural rearrangement or disruption of contact continuity within the candidate regions (Fig. 5 and Figure S3). These results indicate that haplotype-switching artifacts were unlikely to account for the four intervals exceeding 10 cM and support the original haplotype assignments in the reconstructed linkage map.

### HRGM enables QTL detection for root development traits

To further assess the utility of the HRGM-derived linkage map, we quantified eight root-related traits across the S_1_ population, which showed approximately normal distributions despite the relatively small population size (Figure S4). Genome-wide QTL scans were performed for each trait using trait-specific permutation thresholds (Figure S5). Two significant QTLs associated with R-score were detected on chromosomes 1 and 5, whereas no genome-wide significant QTLs were detected for the other seven traits (Fig. 4 and Table S3). qRS1.1 on chromosome 1 spanned 107.0–126.5 cM (17.41–19.14 Mb), with a LOD score of 3.20 and an estimated phenotypic variance explained (PVE) of 26.4%. qRS5.1 on chromosome 5 spanned 44.6–58.1 cM (13.39–14.44 Mb), with a LOD score of 4.16 and an estimated PVE of 32.9%. The higher LOD score and PVE of qRS5.1 indicated a stronger association with R-score than qRS1.1. Together, these results demonstrate that the HRGM-derived map supports QTL detection and trait dissection in a heterozygous species.

## Discussion

The rapid expansion of multiple chromosome-scale genome assemblies has outpaced the development of independent validation frameworks (Shirasawa *et al*., 2021; Song *et al*., 2023). Although long-read sequencing and Hi-C enable highly contiguous assemblies, these approaches primarily enforce structural consistency rather than biological correctness (Rhie *et al*., 2020; Yamaguchi *et al*., 2021). As a result, distinguishing true structural variation from assembly artifacts remains a persistent challenge, particularly in repeat-rich, heterozygous and perennial plant genomes. Genetic linkage maps provide a direct, orthogonal measure of genome structure by capturing meiotic recombination (Fierst, 2015). However, their application has become increasingly limited in precisely those species where genome assemblies are now most rapidly accumulating—such as highly heterozygous crops, polyploid crops, perennial species, clonally propagated crops, and wild germplasm resources (Bourke *et al*., 2018; Ceballos *et al*., 2015; Chen *et al*., 2021). This creates a structural imbalance in which genome assembly has become routine, but recombination-based validation remains inaccessible for many non-model plant species (Whibley *et al*., 2021). HRGM addresses this gap by restoring linkage mapping as a practical validation tool without requiring inbred populations. The near-complete marker order concordance observed across all 11 linkage groups in this study demonstrates that even in a heterozygous, non-model species with no prior genetic resources, recombination-based validation is achievable with HRGM (Fig. 4).

Haplotype-resolved assemblies fundamentally alter how genetic variation can be represented and utilized (Zhang *et al*., 2021). Rather than collapsing heterozygosity into a consensus sequence, phased genomes explicitly encode the two parental chromosome sets within a single individual (Guk *et al*., 2022; Kong *et al*., 2023). This effectively transforms heterozygosity from a complication into an informative signal. HRGM leverages this shift by treating haplotype-resolved sequences as phase-known parental references. In this framework, genetic mapping no longer depends on reconstructing a linkage map through controlled crosses but instead derives it directly from the genome assembly itself. As a result, the minimal experimental unit for linkage mapping becomes a single heterozygous individual and its selfed progeny. This represents a conceptual inversion of conventional mapping strategies, in which phase inference is a limiting step. Notably, haplotype phasing has become feasible not only for major crops but also for highly heterozygous and large genomes (Grewal *et al*., 2025; Huang *et al*., 2024; Jang *et al*., 2024), broadening the scope of HRGM to species that have long been excluded from conventional linkage mapping due to their genetic complexity.

By aligning recombination patterns in S_1_ progeny with haplotype structure, HRGM unifies three processes that are typically treated independently: genome assembly validation, linkage mapping, and QTL analysis (Fierst, 2015; Yu *et al*., 2019). The strong concordance between genetic and physical maps observed here indicates that phased assemblies can serve not only as sequence references but also as scaffolds for interpreting recombination landscapes (Fig. 4). Synteny analysis revealed only minimal structural differences between hap1 and hap2 (Figure S2), and the absence of concentrated segregation distortion further supports that the recombination suppression observed around putative centromeric and pericentromeric regions reflects centromeric architecture rather than pervasive structural disruption between haplotypes (Fig. 4B). Together, these results demonstrate that HRGM can separate linkage phase from a single heterozygote, enabling precise recombination inference and simultaneous assembly validation.

This integration has practical implications for trait dissection. In contrast to population-based approaches such as genome-wide association studies, which are sensitive to allelic heterogeneity and population structure (Korte and Farlow, 2013; Tibbs Cortes *et al*., 2021; Zhou *et al*., 2022), HRGM evaluates the segregation of two defined haplotypes within a controlled genetic background. This enables more direct interpretation of genotype–phenotype relationships and facilitates the identification of robust QTL intervals. As such, HRGM provides a complementary framework for validating candidate loci identified through pangenome or GWAS analyses.

Many plant species with high agronomic or medicinal value remain genetically underexplored due to practical constraints on breeding and population development (Chapman *et al*., 2022; Talabi *et al*., 2022). Long generation times, self-incompatibility, and inbreeding depression limit the feasibility of conventional mapping populations, restricting the application of recombination-based genetic analysis (Chen *et al*., 2021; Schoen and Baldwin, 2023; Song *et al*., 2025). HRGM lowers these barriers by enabling linkage mapping in systems where only a single heterozygous individual and its progeny are available. This expands the scope of genetic analysis to a broader range of species, including crop wild relatives, forest trees, and medicinal plants. In these contexts, HRGM can facilitate the development of genetic resources, support domestication efforts, and enable marker-assisted selection in species that have historically been inaccessible to molecular breeding.

Despite its advantages, HRGM depends on the accuracy of haplotype-resolved genome assemblies. Errors in phasing or structural misassemblies can propagate into incorrect marker assignment and linkage inference. In addition, S_1_ populations may exhibit segregation distortion due to deleterious alleles or inbreeding effects, which can influence marker distribution and recombination estimates. These factors highlight the importance of integrating quality control steps for both genome assembly and genotype data. The applicability of HRGM may also vary across biological systems. While the approach is well suited to diploid species capable of selfing or clonal propagation, its extension to highly polyploid genomes or obligate outcrossing species may require methodological adaptation. Future work should evaluate the robustness of HRGM across varying levels of heterozygosity, trait architecture, genome complexity, structural variation, and ploidy. Further development of HRGM could incorporate improvements in long-read sequencing, phasing algorithms, and graph-based genome representations. By eliminating the need for biparental crosses, HRGM enables trait mapping from a single donor parent and supports rapid development of trait-linked markers for targeted marker-assisted selection.

## Experimental procedures

### Plant materials and phenotyping

The CW accession 9p, the same accession used to generate the HiFi and Omni-C sequencing datasets analyzed in this study, was self-pollinated under screenhouse conditions. Seeds obtained from the self-pollinated plants were used to establish the S_1_ mapping population. A total of 52 S_1_ progeny were cultivated under field conditions at Seoul National University Suwon Farm for approximately 10 months and harvested for phenotypic evaluation.

Eight traits related to root development were measured, including fresh weight, root length, longest root length, root thickness, number of primary roots, number of secondary roots, total number of roots and R-score (Figure S4). Because the irregular shape of CW storage roots makes it difficult to objectively represent root thickness using a single diameter measurement, R-score was calculated as √[root weight / (π × root length × root number)] based on a previous study (Yang *et al*., 2025) and used as a phenotypic trait.

### Haplotype-resolved phasing

The HiFi long-read sequencing data were downloaded from NCBI SRA database (SRR34270040). Haplotype-resolved phasing was performed based on the unitigs generated by hifiasm (Cheng *et al*., 2021), following the approach outlined by Feng et al. (Feng *et al*., 2025). The unitigs were phased using the open-source pipeline available at https://github.com/jia-wu-feng/Pan_Bulbosum with slight modifications. A modified assembly pipeline (https://doi.org/10.6084/m9.figshare.33224118) was used for haplotype phasing. Unitigs from homozygous regions, where both haplotypes are identical in sequence, were identified based on read coverage and subsequently excluded from downstream analysis.

### Marker selection

A CW reference genome was downloaded from Figshare (https://doi.org/10.6084/m9.figshare.29399717). Omni-C sequencing data generated from the same CW individual were obtained from the NCBI SRA database (SRR34348259). A total of 52 S_1_ lines were sequenced using the Illumina NovaSeq 6000 platform. The raw reads were quality-trimmed using cutadapt v4.6 (Martin, 2011) and then were mapped to the reference genome using bwa v0.7.17 (Li, 2013). The SNP calling was conducted using bcftools v1.18 with default parameters (Danecek *et al*., 2021). SNP markers for genetic map construction were selected via a custom script (https://doi.org/10.6084/m9.figshare.33224160). This script performs a multi-step selection to identify haplotype-diagnostic loci and assign genotypes to S_1_ individuals. The SNPs were extracted from loci that were homozygous within each haplotype and fixed for opposite alleles between hap1 and hap2. For validation of candidate haplotype-diagnostic SNPs, haplotype-specific unitigs were directly aligned to the reference genome using minimap2 with the asm10 preset. Candidate SNPs were retained only when supported by high-quality, unambiguous alignments from both haplotypes (MAPQ ≥20, identity ≥98%, aligned length ≥1 kb, and best-chromosome alignment fraction ≥0.70).

The SNPs were then filtered based on progeny genotype quality, read depth, and call rate. For individual genotype calls, minimum read depths (DP) of 4 and 6 were applied to homozygous (A or B) and heterozygous (X) genotypes, respectively. Genotype calls below these depth thresholds were coded as missing. At the marker level, loci with a mean genotype quality (mean GQ) ≥ 30, a mean read depth (mean DP) ≥ 5, and a genotype call rate ≥ 80% across the S_1_ progeny were retained. Progeny genotypes were encoded as homozygous for hap1 (A), homozygous for hap2 (B), or heterozygous (X). Markers with multiallelic or erroneous calls were excluded. The filtered markers were retained only if their segregation among S_1_ individuals conformed to the expected 1:2:1 Mendelian ratio (p ≥ 0.001). Following this filtering pipeline, the selected markers were consolidated into bin markers with a minimum crossover interval ratio of 0.01 and a minimum bin size of 1,000 bp using SNPbinner (Gonda *et al*., 2019).

### Genetic map construction and QTL mapping

The linkage groups (LGs) were constructed using the R/ASMap library (Taylor and Butler, 2017). In R/ASMap library, the mstmap function for F_2_ population was applied using the Kosambi mapping function (dist.fun = “kosambi”) and a threshold of p.value = 1 × 10⁻⁶ (Taylor and Butler, 2017). Only LGs containing more than five bin markers were retained. QTL mapping was performed using the multiple-QTL model mapping (MQM) procedure implemented in the R/qtl package (Arends *et al*., 2010). Genome-wide MQM scans were conducted under a dominance model using a 15-cM window and a 1-cM scanning interval. Trait-specific genome-wide LOD thresholds were determined from 1,000 permutations, and QTLs exceeding the threshold at α = 0.10 were considered significant. QTL support intervals were estimated using a 1.5-LOD drop from the corresponding QTL peak (Arends *et al*., 2010). The synteny blocks were identified using SyRI (Goel *et al*., 2019).

## Supporting Information

Figure S1. Chromosome-wide distribution of informative markers and heterozygosity density.

Figure S2. Macro synteny between haplotype 1 and haplotype 2 genome assemblies.

Figure S3. Evaluation of haplotype-phase consistency at three additional regions using A/B genotype-state swap tests.

Figure S4. Phenotypic distribution of eight traits related to root development in the S_1_ population.

Figure S5. QTL mapping for eight root traits in C. wilfordii.

Table S1. Statistics of haplotype-phased unitigs of the S_0_ individual.

Table S2. Basic information of the genetic map.

Table S3. Summary of the final refined QTLs associated with root development.

## Declarations

### Ethics approval and consent to participate

Not applicable.

### Consent for publication

Not applicable.

### Availability of data and materials

The raw sequencing reads from the S_1_ population were deposited in the NCBI Sequence Read Archive under BioProject accession PRJNA1446679. The haplotype-phased unitig sequences and artificial 150-mer reads are available in Figshare (doi: 10.6084/m9.figshare.31926555). The custom code used for haplotype-diagnostic marker selection and progeny genotyping is archived in Figshare (doi: 10.6084/m9.figshare.33224160) and is also available in the GitHub repository (https://github.com/insulim1014/haplotype-resolved_genetic_mapping). The modified haplotype-phasing assembly code is deposited in Figshare (doi: 10.6084/m9.figshare.33224118) and is also accessible through GitHub (https://github.com/whyeonc/C.wilfordii_haplotype_phasing_code).

### Author Contributions

**Insu Lim**: Conceptualization, Data curation, Formal analysis, Methodology, Visualization, Writing – original draft. **Sae Hyun Lee**: Conceptualization, Methodology, Resources, Writing – original draft. **Woohyeon Cho**: Data curation. **Junhyun Kim**: Resources. **Jee Young Park**: Resources, Supervision. **Murukarthick Jayakodi**: Supervision, Writing – review & editing. **Tae-jin Yang**: Supervision, Writing – review & editing.

## Acknowledgements

The haplotype-phased assembly code was generously contributed by Jia-Wu Feng and Martin Mascher of Leibniz Institute of Plant Genetics and Crop Plant Research (IPK).

