## Supporting Information for "Haplotype-resolved genetic mapping using selfed progeny enables simultaneous genome assembly validation and trait dissection in a heterozygous plant"

**Supplementary Figure 1. Chromosome-wide distribution of informative markers and heterozygosity density.** For each of the 11 chromosomes, informative marker positions are plotted along each chromosome. Above each chromosome, heterozygosity density is shown as the number of informative variant sites per megabase using a 20 kb window size.

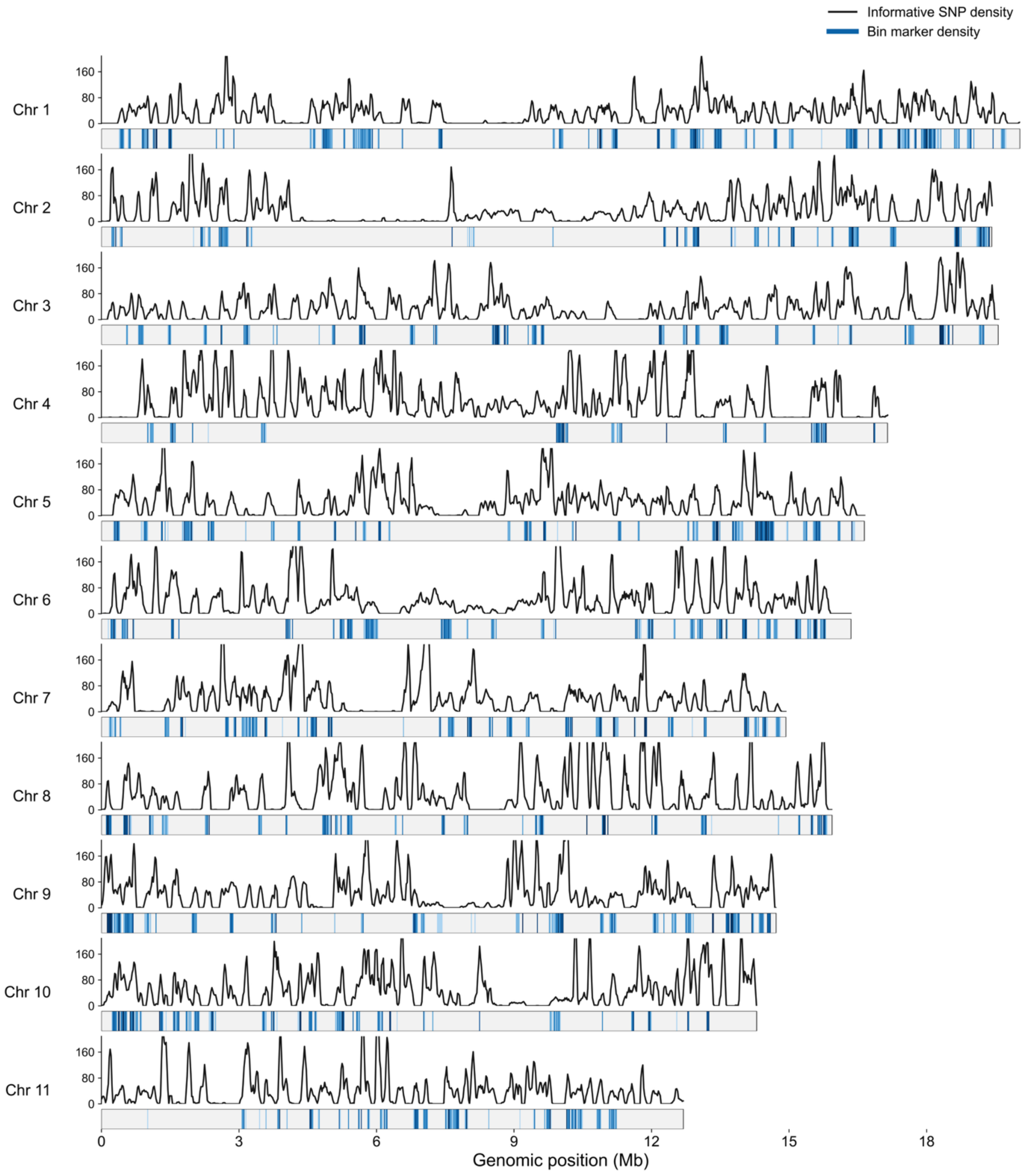

**Supplementary Figure 2. Macro synteny between haplotype 1 and haplotype 2 genome assemblies.**

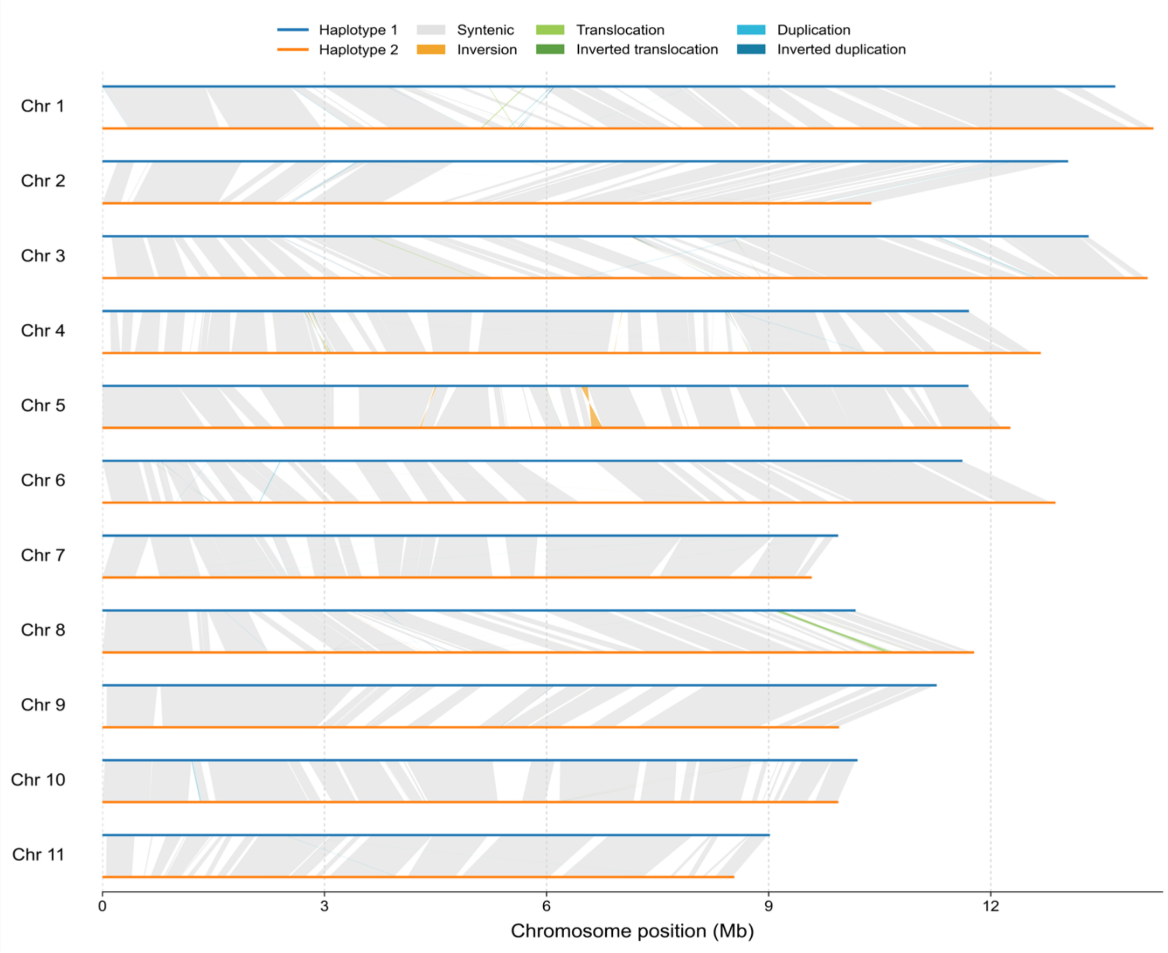

**Supplementary Figure 3. Evaluation of haplotype-phase consistency at three additional regions using A/B genotype-state swap tests. (A)** Comparison of the original and reconstructed linkage maps for target bins chr1-17200461, chr5-14529016, and chr10-3788255, at which abrupt increases in genetic distance exceeding 10 cM were observed. For each region, the left panels show the original genetic map and $S_1$ bin-genotype pattern, whereas the right panels show those reconstructed after reciprocal exchange of the A and B genotype states of the target bin. **(B)** Local Omni-C contact maps of the three target regions. Orange dashed lines indicate the boundaries of each target bin, and the color scales represent the log-transformed Omni-C contact signal.

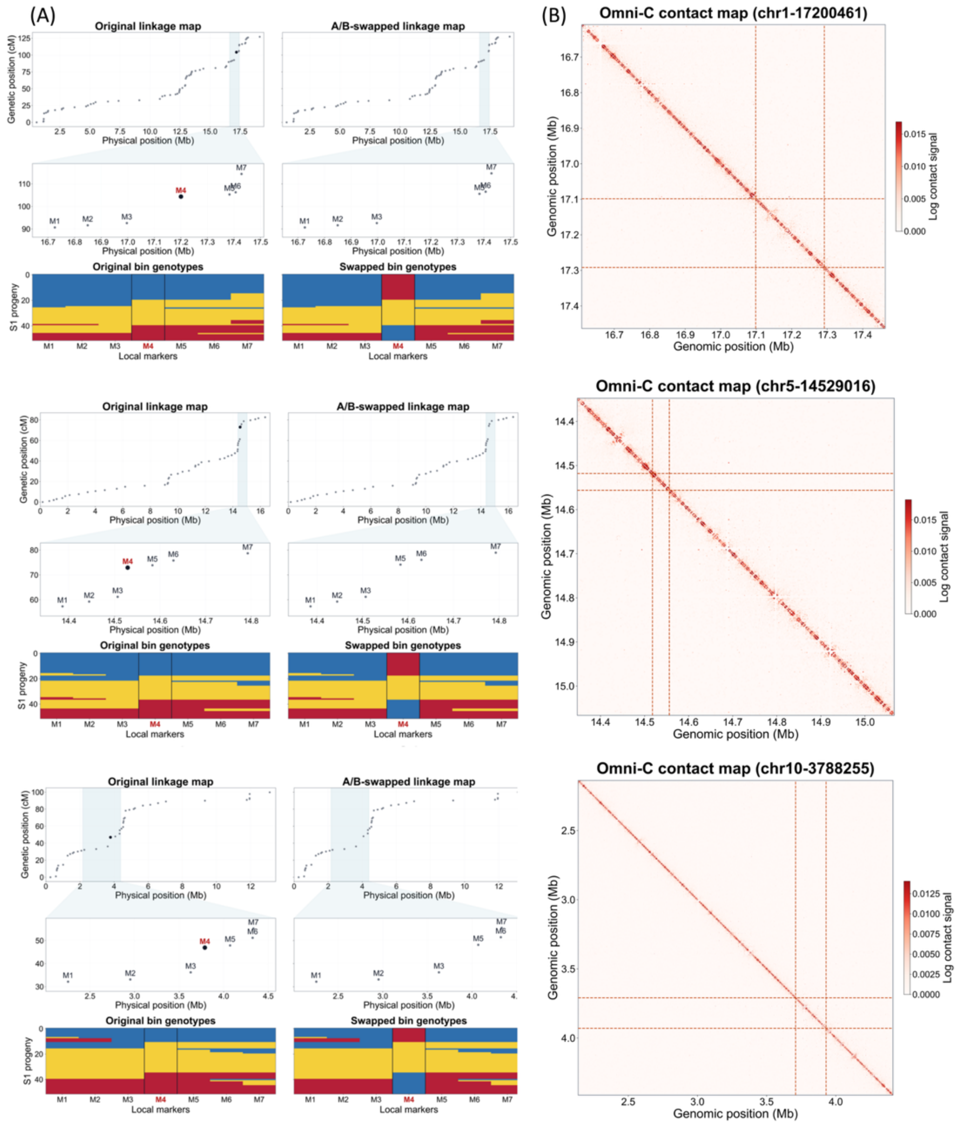

**Supplementary Figure 4. Phenotypic distribution of eight traits related to root development in the $S_1$ population.**

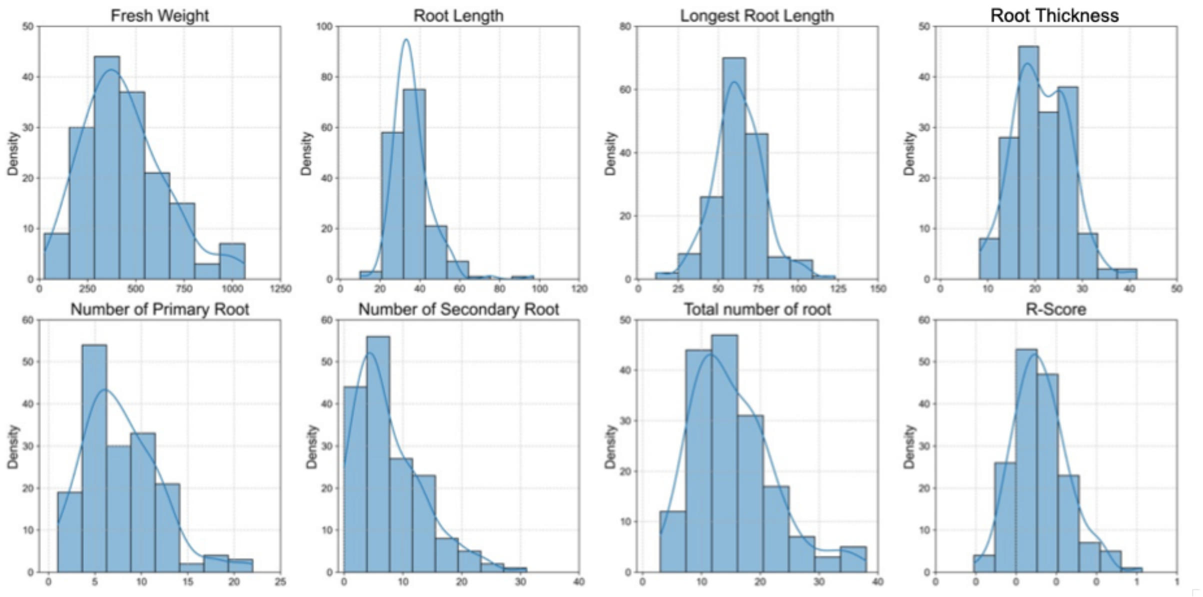

**Supplementary Figure 5. QTL mapping for eight root traits in *C. wilfordii*.** Genome-wide multiple-QTL model mapping (MQM) LOD profiles are shown for each trait across the 11 linkage groups. The red dotted line in each panel indicates the trait-specific genome-wide LOD threshold determined by 1,000 permutations at α = 0.10.

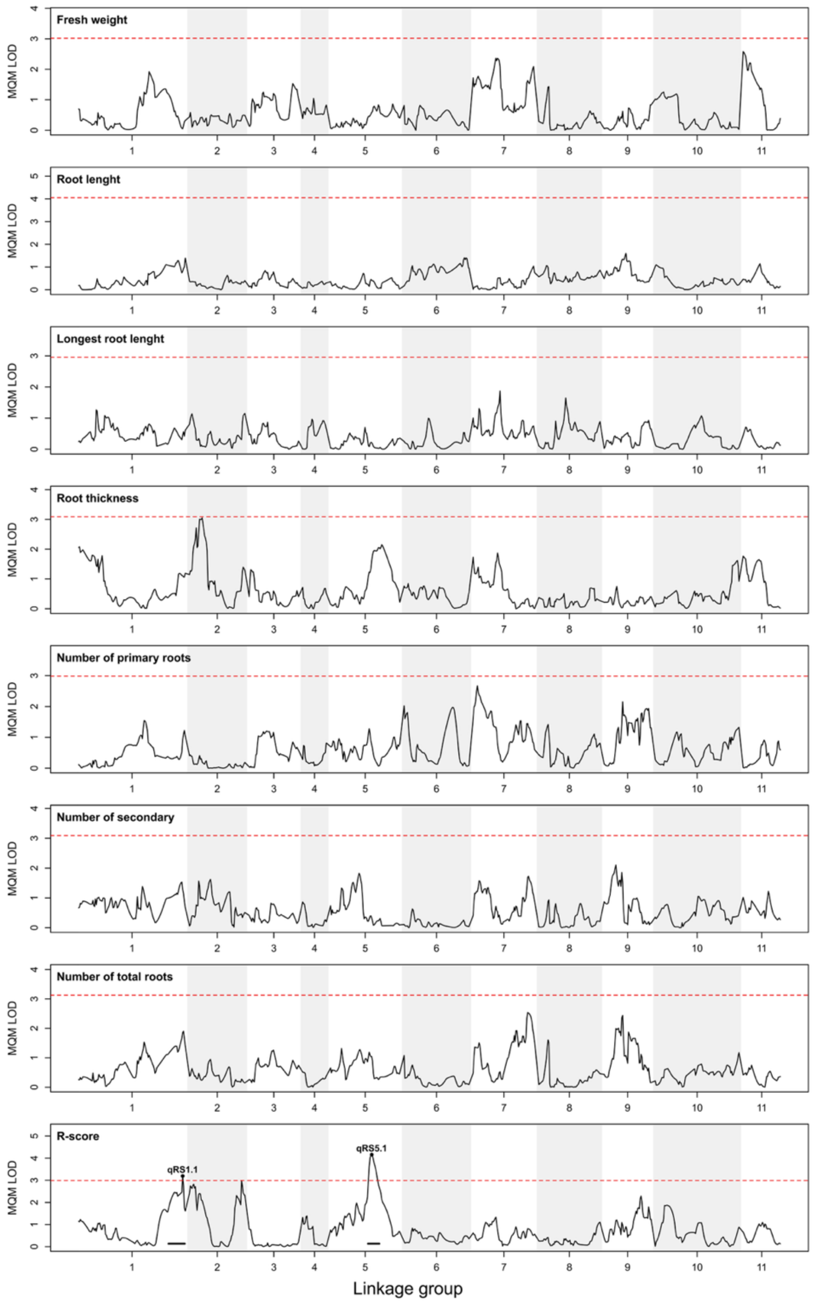

**Supplementary Table 1. Statistics of haplotype-phased unitigs of the $S_0$ individual.**

| Chromosome | Total unitig size (Mb) | | | Number of unitigs (*n*) | | |
|---|---|---|---|---|---|---|
| | Unphased | Haplotype 1 | Haplotype 2 | Unphased | Haplotype 1 | Haplotype 2 |
| Chr 1 | 0.1 | 13.7 | 14.3 | 1 | 72 | 85 |
| Chr 2 | 2.4 | 13 | 10.4 | 6 | 67 | 56 |
| Chr 3 | 0 | 13.4 | 14.1 | 0 | 72 | 74 |
| Chr 4 | 0 | 11.7 | 12.7 | 0 | 52 | 53 |
| Chr 5 | 0 | 11.7 | 12.2 | 0 | 61 | 63 |
| Chr 6 | 0 | 11.6 | 12.8 | 0 | 57 | 66 |
| Chr 7 | 0 | 10.1 | 9.6 | 0 | 59 | 60 |
| Chr 8 | 0 | 10.2 | 12 | 0 | 61 | 66 |
| Chr 9 | 0 | 11.2 | 9.9 | 0 | 51 | 53 |
| Chr 10 | 0 | 10.2 | 9.9 | 0 | 55 | 54 |
| Chr 11 | 0 | 9 | 8.5 | 0 | 50 | 45 |
| Total | 2.5 | 125.7 | 126.5 | 7 | 657 | 675 |

**Supplementary Table 2. Basic information of the genetic map.**

| Group | Total length (cM) | No. of bin markers | Physical length (Mb) | Mean inter-marker interval (cM) |
|---|---|---|---|---|
| LG1 | 127.15 | 68 | 20.04 | 1.90 |
| LG2 | 66.09 | 45 | 19.42 | 1.50 |
| LG3 | 58.85 | 37 | 19.56 | 1.64 |
| LG4 | 28.21 | 21 | 17.15 | 1.41 |
| LG5 | 82.57 | 47 | 16.64 | 1.80 |
| LG6 | 77.42 | 36 | 16.35 | 2.21 |
| LG7 | 73.27 | 52 | 14.93 | 1.44 |
| LG8 | 72.85 | 46 | 15.94 | 1.62 |
| LG9 | 55.92 | 45 | 14.72 | 1.27 |
| LG10 | 99.79 | 41 | 14.29 | 2.50 |
| LG11 | 44.30 | 26 | 12.69 | 1.77 |
| **Total** | **786.4** | **464** | **181.73** | **1.74** |

**Supplementary Table 3. Summary of the final refined QTLs associated with root development.**

| Trait | QTL | Chr. No. | 1.5-LOD support interval (cM) | Physical interval (bp) | LOD | PVE (%) | P-val |
|---|---|---|---|---|---|---|---|
| R-score† | qRS1.1 | 1 | 107–126.5 | 17,406,373–19,137,184 | 3.20 | 26.4 | 0.0629 |
| R-score† | qRS5.1 | 5 | 44.6–58.1 | 13,394,486–14,444,373 | 4.16 | 32.9 | 0.013 |

†R-score means a normalized value of root thickness for measuring root with irregular shape.
